# LRX- and FER-dependent extracellular sensing coordinates vacuolar size for cytosol homeostasis

**DOI:** 10.1101/231043

**Authors:** Kai Dünser, Shibu Gupta, Christoph Ringli, Jürgen Kleine-Vehn

## Abstract

Cellular elongation requires the defined coordination of intra- and extracellular processes. The vacuole is the biggest plant organelle and its dimension has a role in limiting cell expansion (Löfke et al., 2015; Scheuring et al., 2016). We reveal that the increase in vacuolar occupancy enables cellular elongation with relatively little enlargement of the cytosole. It remains, however, completely unknown how the vacuolar size is coordinated with other growth-relevant processes. Intriguingly, we show that extracellular constraints impact on the intracellular expansion of the vacuole. The underlying cell wall sensing mechanism requires the interaction of the extracellular leucine-rich repeat extensin (LRX) with the receptor-like kinase Feronia (FER). Our data suggests that LRX links the plasma membrane localised FER with the cell wall, allowing this module to jointly sense and convey extracellular signals to the underlying cell. This mechanism coordinates cell wall acidification/loosening with the increase in vacuolar size, contributing cytosol homeostasis during plant cell expansion.

---

Plant cells are constrained by the surrounding, rigid cell wall. Accordingly, cellular expansion requires a complex coordination of several internal and external processes. Developmentally defined cell wall loosening and concomitant water uptake are important pre-requisites to enable turgor-driven cell expansion (reviewed in Braidwood et al., 2013). On the other hand, the dimension of the biggest plant organelle, the vacuole, does not only correlate with cell size in plants, but phytohormone-dependent interference with its expansion also reduces cellular elongation rates (Löfke et al., 2015; Scheuring et al., 2016). Despite its importance, the regulation of vacuolar size and its coordination during cellular elongation is yet an under-investigated research area.

Here we use epidermal atrichoblast (non-root hair forming) root cells to record vacuole expansion during cellular elongation. To assess the relative dilation of the vacuole, we combined the fluorescent dye BCECF (2’,7’-Bis-(2-Carboxyethyl)-5-(and-6)-Carboxyfluorescein), which accumulates in the vacuolar lumen of plant cells, with propidium iodide which stains the exterior of the cells (Scheuring et al., 2015). Subsequently, we performed defined z-stack imaging and 3D rendering of the cell, allowing quantification of the vacuolar occupancy of the cell (Scheuring et al., 2016). In meristematic cells, the vacuole occupied about 30-40% of the cellular space (Figure 1 a and b). By contrast, the size of the vacuole dramatically increased during cellular elongation, ultimately taking up 80-90% of the cellular volume (Figure 1 a and b). While the epidermal cells enlarged their volume by a factor of 14, the absolute space between the vacuole and the cell boundary increased only about 3-fold (Figure 1 c and Suppl. Figure 1 a and b). The 3D imaging of a cytosolic fluorescent protein also confirmed that the cytosol shows limited volume expansion during epidermal elongation (Suppl. Figure 1 a and b). Accordingly, the relative increase in vacuolar size has a dramatic impact on cytosol homeostasis during cellular enlargement. We conclude that the size of the vacuole is important for rapid plant cell expansion, requiring relatively little *de-novo* production of cytosolic components.

**Figure 1.**
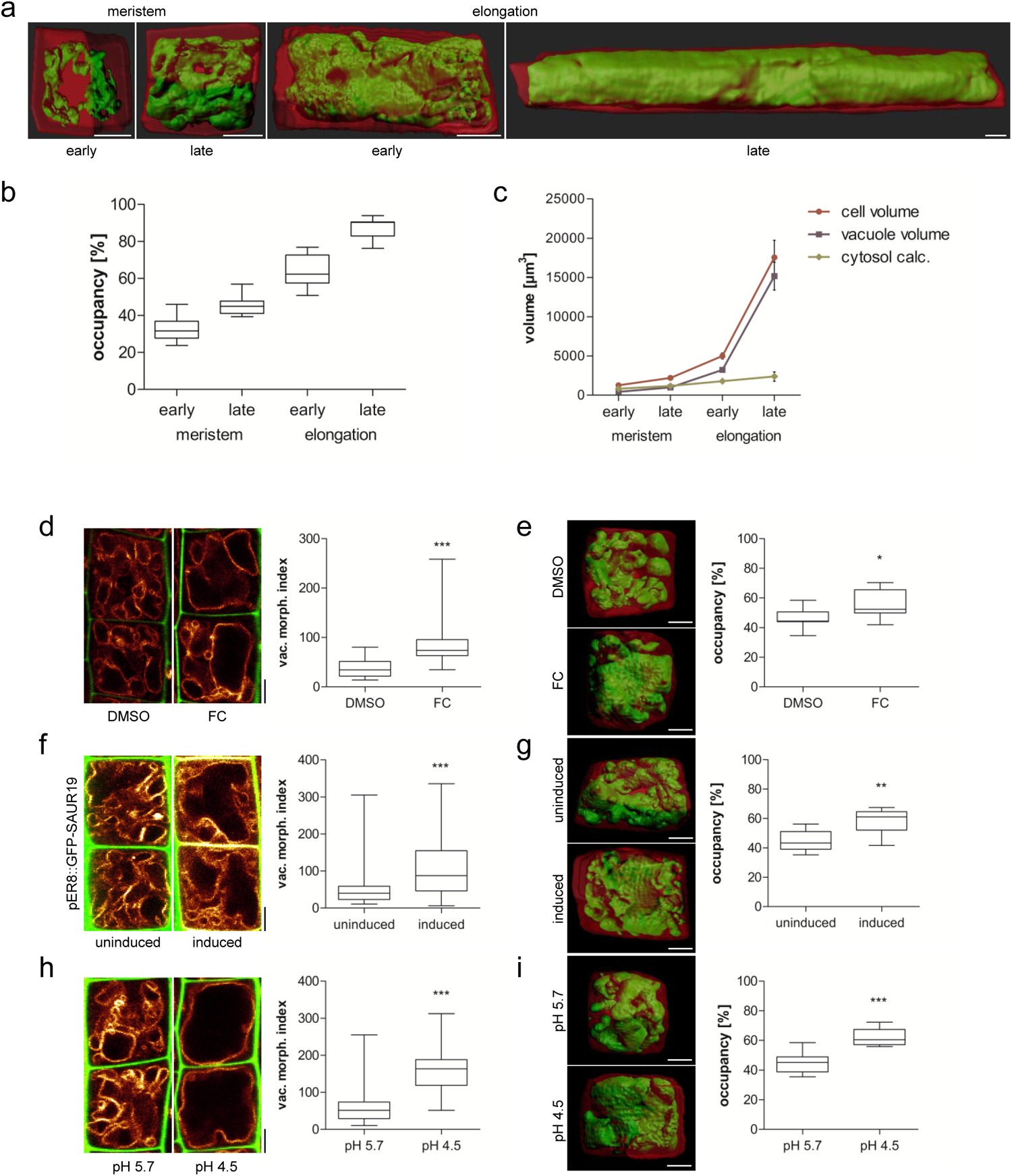
Vacuolar size correlates with cell wall acidification/loosening. **a)** 3-D reconstructions of propidium iodide (PI)-stained cell walls (red) and BCECF-stained vacuoles (green) of epidermal atrichoblasts in the early and late meristem and in the early and late elongation zone. **b)** Boxplots showing vacuolar occupancy of cells in the defined zones (n=7-11). **c)** Graph depicts absolute values of cell volume and vacuolar volume in the depicted root zones (data points display mean, error bars depict s.e.m.). “Cytosol” is approximated as the difference between cell volume and vacuolar volume. **d)** Representative images and quantification of vacuolar morphology of late meristematic cells. PI (green) and pUBQ10::VAMP711 (yellow) depict cell wall and tonoplast, respectively. Seedlings were treated with DMSO (solvent control) or 5 μM FC for 2.5 h in liquid medium (n=24). **e, g, h)** 3-D reconstructions of PI-stained cell wall (red) and BCECF-stained vacuole (green) of late meristematic cells. **e)** Boxplot depicts vacuolar occupancy of the cell treated with the solvent DMSO or 5 μM FC for 2.5 h in liquid medium (n = 11). **f, g)** Cell wall and vacuolar membrane was visualized with PI (green) and MDY-64 (yellow) **(f)** or pUBQ10::VAMP711 **(g)** (yellow), respectively. **f)** pER8::GFP-SAUR19 seedlings were treated with DMSO (n=60) or 10 μM β-estradiol (n=56) for 6 h in liquid medium. **g)** 3-D cell reconstructions of pER8::GFP-SAUR19 lines. Boxplot depicts vacuolar occupancy of the cell. Seedlings were treated with the solvent control DMSO (n=11) or 10 μM β-estradiol (n=8) for 6 h in liquid medium. **h)** Col-0 wild type seedlings were treated for 3 h in liquid medium adjusted to pH 5.7 (n=44) or pH 4.5 (n=40). **i)** Boxplot depicts vacuolar occupancy of cell. Seedlings were treated for 3 h in liquid medium adjusted to pH 5.7 or pH 4.5 (n=11). **a-i)** Scale bars: 5 μm. boxplots: whiskers display min. to max. values. Statistical analyses were performed using Student’s *t*-test, *p ≤ 0.05; **p ≤ 0.01, ***p ≤ 0.001. Representative experiments are shown.

Cell wall acidification is central in activating a cascade of events, ultimately leading to cell wall loosening and subsequent cellular elongation (Fendrych et al., 2016; Barbez et al., 2017). Accordingly, cell wall acidification/loosening (Barbez et al., 2017) correlates with the increase in vacuolar size (Figure 1a-c), but the coordinative mechanism is unknown. We, therefore, investigated the possibility that apoplast acidification/cell wall loosening is sensed and provides a feedback for vacuolar morphogenesis. We either directly acidified the rhizosphere (by lowering the pH of the media) or genetically (SAUR19 induction) as well as pharmacologically (fusicoccin treatment) induced the activity of the plasma membrane H+-ATPase (Spartz et al., 2014; Spartz et al., 2016, Marre, 1979). All these conditions have recently been shown to acidify the pH of epidermal cell walls, consequently inducing cellular elongation (Barbez et al., 2017). We visualized the vacuolar morphology in root epidermal cells in the late meristematic zone, allowing assessment of the impact of cell wall acidification before the onset of elongation. Using the tonoplast stain MDY-64 or the tonoplast marker line pUBQ10::VAMP711-YFP (Scheuring et al., 2015; Löfke et al., 2015), we revealed that cell wall acidification caused a dramatic alteration in vacuolar morphology (Figure 1 d, f, h). Cell wall acidification did not only change the vacuolar appearance, but also induced an increased vacuolar occupancy of the cell (Figure 1 e, g, i). Our data accordingly suggests that cell wall acidification/loosening impacts on vacuolar morphogenesis and size.

The inhibition of pectin methyl esterases (PMEs) leads to reduced cellular elongation, presumably due to stiffening of the cell walls (Wolf et al., 2012). Therefore, we used epigallocatechin gallate (EGCG), which is a natural inhibitor for PMEs (Lewis et al., 2008), to inducibly interfere with the properties of the cell wall. The application of EGCG induced smaller vacuolar structures and reduced the overall vacuolar occupancy of the cell (Figure 2 a and b). Notably, roots that penetrate a relatively stiff medium similarly show smaller vacuoles, when compared to surface-grown roots (Figure 2 c and d). These findings suggest that extracellular constraints restrict intracellular enlargements of vacuoles.

**Figure 2.**
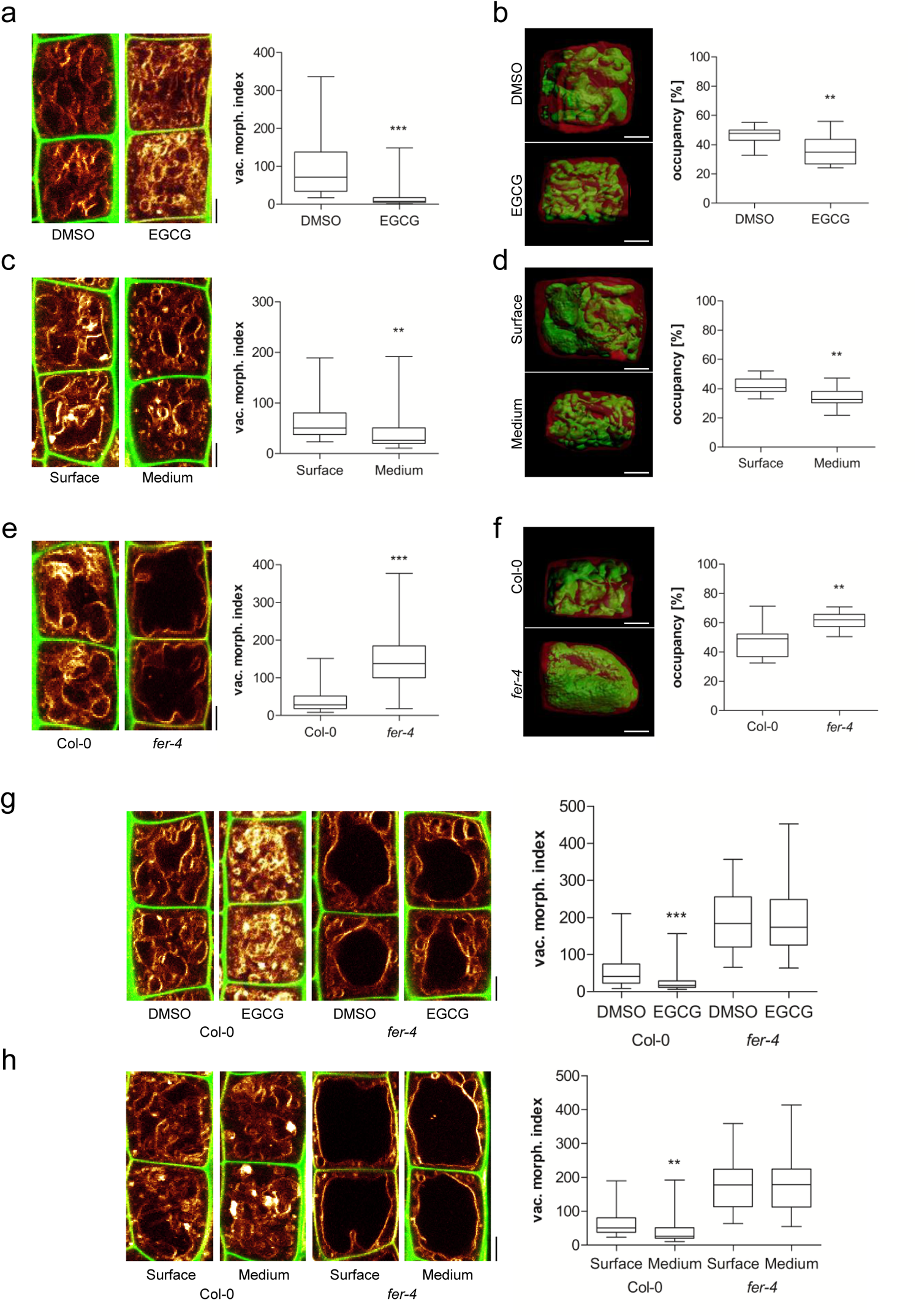
Putative cell wall sensor Feronia impacts on vacuolar size. **a-h)** Representative images and quantification of vacuolar morphology of late meristematic atrichoblast cells. **a, c, e, g, h)** PI (green) and MDY-64 (yellow) staining depicts cell wall and vacuolar membrane, respectively. **b, d, f)** 3-D reconstructions of PI-stained cell wall (red) and BCECF-stained vacuole (green). **a)** Seedlings were treated with DMSO solvent control or 50 μM EGCG for 22 h on solid medium (n=40). **b)** Boxplot depicts vacuolar occupancy of the cell. Seedlings were treated with DMSO (n=16) or 50 μM EGCG (n=15) for 22 h on solid medium. **c)** Seedling roots were grown on the surface (n=40) or into the matrix (n=48) of 2 % agar-containing solid medium. **d)** Boxplot depicts vacuolar occupancy of surface (n=10) and into the medium (n=11) grown seedling roots. **e)** Vacuolar morphology of Col-0 (n=64) and *fer-4* (n=60). **f)** Boxplot depicts vacuolar occupancy of Col-0 control (n=11) and *fer-4* (n=10) mutant cells. **g)** Col-0 (n=52) and *fer-4* (n=44) seedlings were treated with solvent DMSO or 50 μM EGCG for 22 h on solid medium. **h)** Col-0 (n=40-48) and *fer-4* (n=32) mutant seedling roots were grown on the surface or into the matrix of 2 % agar-containing solid medium. **a-h)** Scale bars: 5 μm. Boxplots: whiskers display min. to max. values. Statistical analyses were performed using Student’s *t*-test, *p ≤ 0.05; **p ≤ 0.01, ***p ≤ 0.001. Representative experiments are shown.

We assume that cell wall sensing integrates vacuolar size. To test this hypothesis, we focused on the receptor-like kinase (RLK) FERONIA (FER), which is required for mechanical cell wall sensing (Shih et al., 2014). The *fer-2* and *fer-4* loss-of-function mutants showed enlarged, roundish vacuoles (Figure 2 e; Suppl. Figure 2 a) and furthermore increased the vacuolar occupancy of the epidermal cells (Figure 2 f; Suppl. Figure 2 b). Moreover, *fer* mutant vacuoles were neither affected by EGCG treatments nor by extracellular constraints of the substrate (Figure 2 g and h). Accordingly, we conclude that an extracellular, FER-dependent signal restricts intracellular expansion of the vacuole.

FER is a transmembrane protein, with an extracellular domain for signal perception and a cytoplasmic domain that phosphorylates target molecules (Escobar-Restrepo et al., 2007). Notably, *pFER::FERKR-GFP* bares a mutation in the kinase domain and was not able to complement the vacuolar phenotype of *fer* mutants (Suppl. Figure 2 c and d). This data supports a role for the FER kinase activity in restricting intracellular expansion of the vacuole.

FER is a versatile growth integrator in plants with important contributions to a wide range of responses (Li et al., 2016). It remains, however, still unknown how FER contributes to the mechanical sensing of the extracellular space (Shih et al., 2014). Next, we turned our attention to extracellular proteins with a possible role in cell wall sensing and regulation of vacuolar size. Interestingly, leucine-rich repeat extensins (LRXs) are extracellular proteins (Baumberger et al., 2003) and showed co-expression with FER and other FER-related RLKs (Suppl. Figure 3 a and b). LRXs display an N-terminal leucine-rich repeat (LRR) and a C-terminal extensin (EXT) domain (Draeger et al., 2015). While the LRR domain is presumably involved in protein-protein interactions, the EXT domain allows the LRX proteins to bind cell wall components (Ringli, 2010). This interesting domain structure also envisioned the hypothetical role of LRX in cell wall sensing (Humphrey et al., 2007). Intriguingly, *lrx3 lrx4 lrx5* triple mutants displayed larger vacuolar lumina (Figure 3 a). Similar to *fer* mutants, these changes also resulted in vacuoles occupying more space in the late meristematic, epidermal cells (Figure 3 b). *lrx3 lrx4 lrx5* triple mutant vacuoles were moreover fully resistant to EGCG treatments as well as to external constraints by the substrate (Figure 3 c and d). We accordingly conclude that extracellular LRX proteins are involved in setting the intracellular expansion of the vacuole.

**Figure 3.**
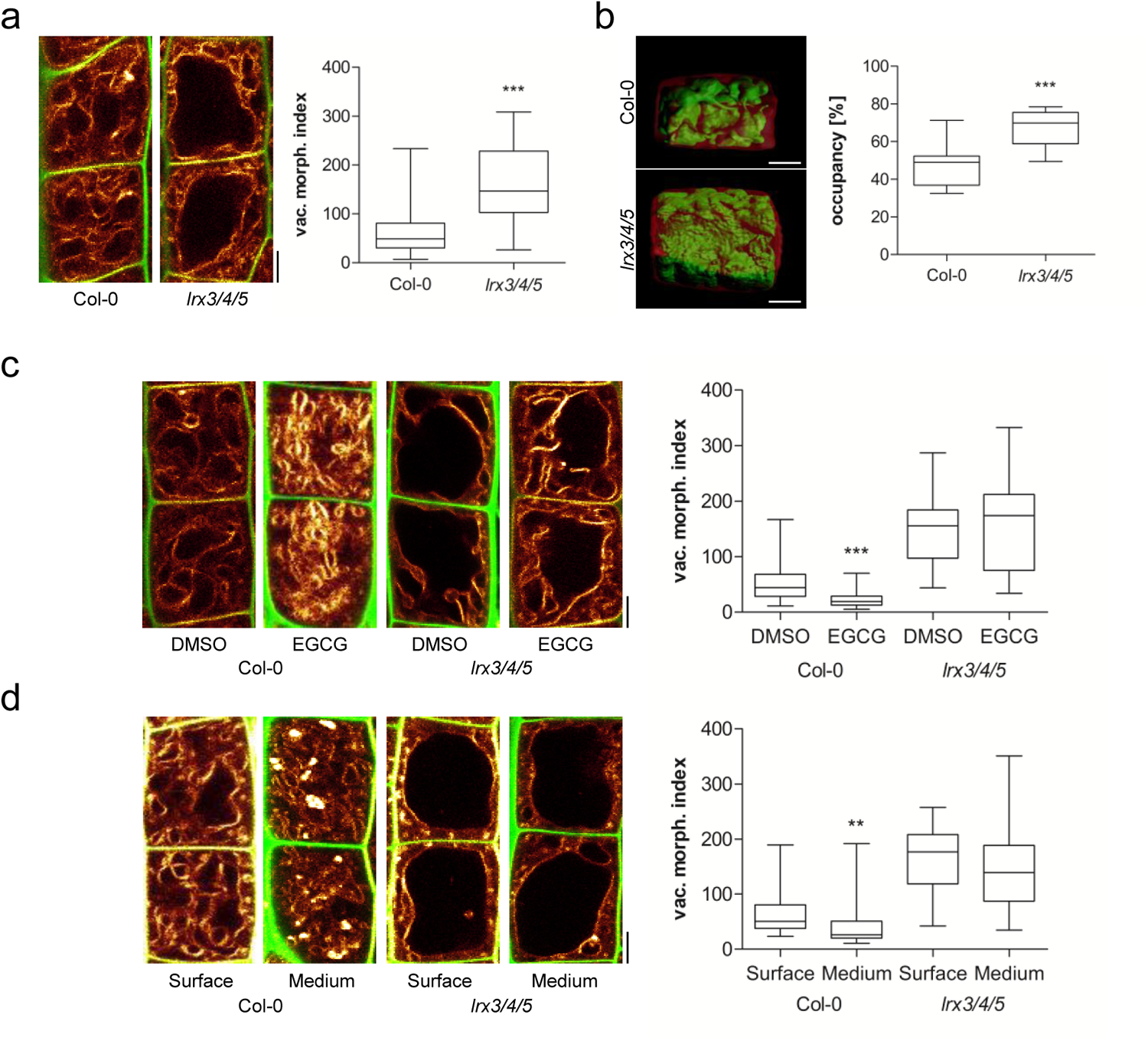
Extracellular LRX proteins are required to constrain vacuolar expansion. **a-d)** Representative images and quantification of vacuolar morphology of late meristematic atrichoblast cells. **a, c, d)** PI (green) and MDY-64 (yellow) staining depicts cell wall and vacuolar membrane, respectively. **b, d, f)** 3-D reconstructions of PI-stained cell wall (red) and BCECF-stained vacuole (green) of late meristematic cells. **a)** Vacuolar morphology of Col-0 control (n=52) and *lrx3/4/5* triple mutants (n=48). **b)** Boxplot depicts vacuolar occupancy of the cell in Col-0 control (n=11) and *lrx3/4/5* (n=10). **c)** Col-0 (n=40-44) and *lrx3/4/5* (n=36) seedlings were treated with DMSO or 50 μM EGCG for 22 h on solid medium. **d)** Col-0 (n=40-48) and *fer-4* (n=28-32) seedlings were grown on the surface or into the matrix of 2 % agar-containing solid medium. **a-d)** Scale bars: 5 μm. Boxplots: whiskers display min. to max. values. Statistical analyses were performed using Student’s *t*-test, *p ≤ 0.05; **p ≤ 0.01, ***p ≤ 0.001. Representative experiments are shown.

Importantly, not only the vacuoles, but also the overall plant phenotype, of *lrx3 lrx4 lrx5* triple mutants closely resembled the appearance of *fer* mutants (Figure 4 a). Moreover, the *fer lrx3 lrx4 lrx5* quadruple mutants were not distinguishable from the *fer* single mutants (Figure 4 a), indicating that FER and LRX function in the same pathway.

**Figure 4.**
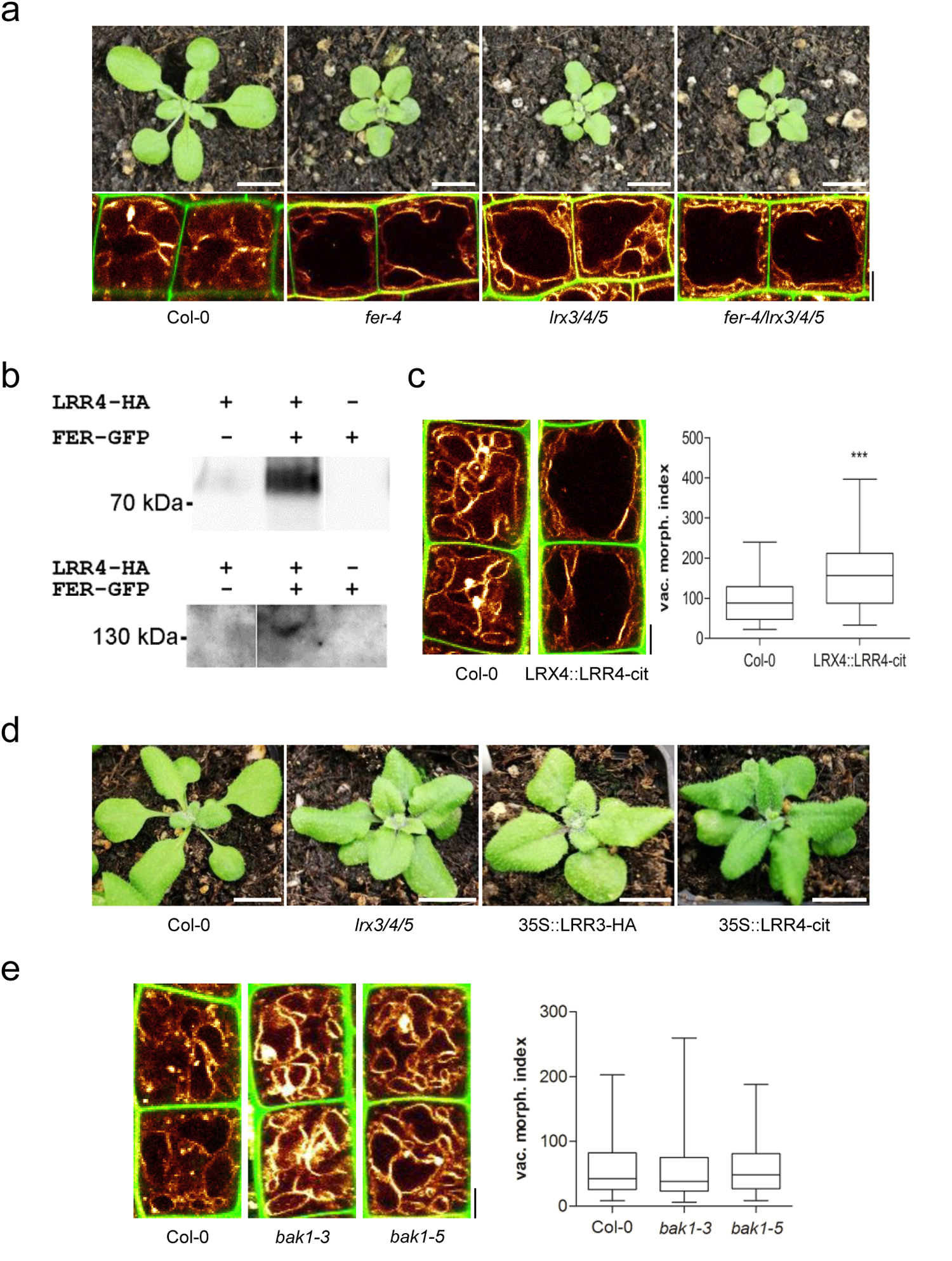
LRX and Feronia jointly sense extracellular signals. **a)** Rosette phenotype of 3-weeks-old Col-0, *fer-4*, *lrx3/4/5* and *fer-4/lrx3/4/5* (upper panel). Vacuolar morphology (MDY-64-stained) of late meristematic atrichoblast cells of Col-0, *fer-4*, *lrx3/4/5* and *fer-4/lrx3/4/5* (lower panel). **b)** LRR4-HA and FER-GFP were transiently expressed (as indicated by + and -). Proteins were either precipitated with anti-GFP (upper panel) or anti-HA (lower panel) antibody followed by immunodetection by Western blotting with an anti-HA antiserum (upper panel) or anti-GFP antiserum (lower panel), respectively. **c)** Representative images and quantification of vacuolar morphology of late meristematic cells of Col-0 and *LRX4::LRR4-Citrine* (n=44). PI (green) and MDY-64 (yellow) staining depict cell wall and tonoplast, respectively. **d)** Rosette phenotype of 3-weeks-old Col-0, *lrx3/4/5*, 35S::LRR3-HA and *35S::LRR4-Citrine*. **e)** Representative images and quantification of vacuolar morphology of late meristematic cells of Col-0, *bak1-3* and *bak1-5* (n=48). PI (green) and MDY-64 (yellow) staining depict cell wall and vacuolar membrane, respectively. **a-e)** Scale bars: 5 μm. Boxplots: whiskers display min. to max. values. Statistical analyses were performed using Student’s *t*-test, *p ≤ 0.05; **p ≤ 0.01, ***p ≤ 0.001. Representative experiments are shown.

Notably, truncated LRX proteins that lack the EXT cell wall binding domain partially associate with the plasma membrane (Ndinyanka et al., 2017), suggesting that the N-terminus of LRX4 interacts with some membrane component. Accordingly, we expressed the HA-tagged N-terminal part of LRX4 (LRR4-HA) together with a GFP-tagged FER in *tobacco*. Co-immunoprecipitation of FER and LRR4 indicated an association of these two proteins in a complex (Figure 4 b).

Most remarkably, we noticed that the expression of a citrine-tagged LRR4 under the LRX4 promoter (pLRX4::LRR4-Citrine) was sufficient to increase the size of vacuoles in late meristematic, epidermal cells (Figure 4 c). This suggests that the EXT cell wall binding domain is critically important to repress the expansion of the vacuole. Moreover, overexpression of LRR4-Citrine and LRR3-HA (similar truncation of LRX3) caused dominant negative phenotypes that were reminiscent of the *lrx3 lrx4 lrx5* triple or *fer* single mutants (Figure 4 d).

This set of data indicates that interaction of LRX and FER contributes to extracellular sensing, impacting on intracellular processes, such as the expansion of the vacuole. In addition to its extracellular collaboration with LRX, FER interacts within the plasma membrane with the co-receptor BRI1-associated kinase 1 (BAK1)/ somatic embryogenesis receptor kinase 3 (SERK3) (Stegmann et al., 2017). This FER/BAK1 complex functions as a scaffold, allowing the recruitment of other RLKs, such as immune response receptors EF-TU RECEPTOR (EFR) and FLAGELLIN-SENSING 2 (FLS2) (Stegmann et al., 2017). Therefore, we analysed the contribution of the BAK1 co-receptor to the regulation of vacuolar form. Importantly, none of the tested *bak1* (strong and weak) mutant alleles affected the vacuolar morphology (Figure 4 e). Accordingly, we conclude that the FER/LRX-dependent cell wall sensing mechanism is distinct from the previously reported BAK1-dependent scaffold mechanism.

The increase in vacuolar volume allows for rapid cell elongation with relatively little de-novo production of cytosolic content. Our data suggests that a cell wall sensing mechanism impacts on the intracellular expansion of the vacuole. This feedback ensures the spatial and temporal coordination of cell wall acidification/loosening with the increase in vacuolar size, which effects cytosol homeostasis during rapid cell expansion.

FER has been previously proposed to be required for cell wall sensing (Shih et al., 2014), but the underlying mechanism remains unknown. Our work proposes LRX as a physical link between the plasma membrane localised FER on one side (via the LRR domain) and the cell wall (via the EXT domain) on the other. Both LRX and FER are essential for constraining the vacuole, suggesting that the interaction of LRX and FER is required for its signalling to the underlying cell. The FER-/LRX-dependent mechanism is independent of the co-receptor BAK1. Therefore, the LRX-dependent cell wall sensing mechanism differs molecularly from the FER scaffold function during immune responses (Stegmann et al., 2017). Accordingly, FER appears to form distinct complexes, which is consistent with the very diverse external signals that can be perceived by the same core component FER. The dominant negative impact of LRR4 suggests the presence of additional components in the FER/LRX complex, assuming that the truncated LRX outcompetes/sequesters some crucial interactor(s). Given the molecular complexity of the FER signalling, it is furthermore likely that the state of LRX/FER interaction also disrupts and/or enables other FER involved interactions (e.g. BAK1).

We propose that an LRX- and FER-dependent module integrates the cell wall status with intracellular expansion of the vacuole. It remains to be seen precisely how LRX/FER signalling at the cell surface leads to the modulation of vacuoles. FER-dependent phosphorylation events impact on RAC/ROP GTPases (Rho-related molecular switches in plants), which are known regulators of actin dynamics (Duan et al., 2010). Notably, the actin cytoskeleton surrounds the vacuole and contributes to the regulation of vacuolar size (Scheuring et al., 2016). Accordingly, it is very tempting to speculate that an actin-dependent process could link the LRX/FER-dependent, extracellular sensing mechanism with the intracellular control of vacuolar expansion.

## Material and methods

### Plant material and growth conditions

Most experiments were carried out in A. thaliana (Col-0ecotype). The following plant lines were described previously: fer-4 (Shih et al., 2014), *fer-2* (Deslauriers and Larsen, 2010), *lrx3/lrx4/lrx5* (Draeger et al., 2015), *pUBQ10::VAMP711-YFP* (Geldner et al., 2009), *pER8::GFP-SAUR19* (Spartz et al., 2014), *pHusion* (Gjetting et al., 2012), *pFER::FERKR-GFP* (Shih et al., 2014) and *LRX4::LRR4-citrine* (Ndinyanka et al., 2017). The *fer-4/lrx3/lrx4/lrx5* quadruple mutant was generated by crossing *lrx3/lrx4/lrx5* (female) with *fer-4* (male). *35S::LRR3-HA* and *35S::LRR4-Citrine* (Ndinyanka et al., 2017) plants were obtained by floral dipping and subsequent selection on Kanamycin. Seeds were stratified at 4°C for 2 days in the dark and were grown on vertically orientated ½ Murashige and Skoog (MS) medium plates under a long-day regime (16 h light/8 h dark) at 20–22°C.

### Chemicals

All chemicals were dissolved in dimethyl sulfoxide (DMSO) and applied in solid or liquid ½ MS medium. MDY-64 was obtained from life technologies (CA, USA), *β*-estradiol, propidium iodide (PI), 2’,7’ -Bis(2-carboxyethyl)-5(6)-carboxyfluorescein acetoxymethyl ester (BCECF-AM) and epigallocatechin gallate (EGCG) from Sigma (MO, USA), fusicoccin (FC) from Cayman Chemical (MI, USA).

### Phenotype analysis

Vacuolar morphology index and occupancy were quantified in 6-d-old seedlings. Confocal images were analyzed using ImageJ (vacuolar morphology index) or processed using Imaris (vacuolar occupancy of cells). To calculate the vacuolar morphology index, the longest and widest distance of the biggest luminal structure was measured and multiplied (Löfke et al., 2015). The atrichoblast cells were quantified before the onset of elongation (late meristematic). To depict this region, the first cell being twice as long as wide was considered as the onset of elongation. Starting from this cell, the next cell towards the meristem was excluded (as it usually shows either partial elongation and/or already substantial vacuolar expansion), and vacuoles of the subsequent 4 cells were quantified as described previously (Scheuring et al., 2016). For the analysis of occupancy, 1 cell in this region was used. Vacuolar shape/size were quantified in at least 8 roots (unless stated otherwise in the figure legend). MDY-64 and BCECF staining was performed as described previously (Scheuring et al., 2015). Plant rosette phenotype evaluation was performed 3 weeks after germination.

### 3-D reconstruction of vacuoles

Imaris 8.4.0 was used for the reconstruction of cell and vacuole volumes. Based on the PI channel, every 3rd slice of the z-stack was utilised to define the cell borders using the isoline, magic wand or manual (distance) drawing functions in the manual surface creation tool. After creating the surface corresponding to the entire cell, a masked channel (based on BCECF) was generated by setting the voxels outside the surface to 0. Subsequently, a second surface (based on the masked BCECF channel) was generated automatically with the smooth option checked. The obtained surface was visually compared to the underlying BCECF channel and, if necessary, the surface was fitted to the underlying signal by adjusting the absolute intensity threshold slider. Finally, volumes of both surfaces were extracted from the statistics window.

### Confocal Microscopy

For image acquisition a Leica TCS SP5 (DM6000 CS) confocal laser-scanning microscope, equipped with a Leica HCX PL APO CS 63 × 1.20 water-immersion objective, was used.MDY-64 was excited at 458 nm (fluorescence emission: 465–550 nm), GFP and BCECF at 488 nm (fluorescence emission: 500–550 nm), YFP at 514 nm (fluorescence emission: 525–578 nm), and PI at 561 nm (fluorescence emission: 644–753 nm). Roots were mounted in PI solution (0.02 mgWe are grateful to Y. Belkhadir/mL) to counterstain cell walls. Z-stacks were recorded with a step size of 420 nm.

### Co-Immunoprecipitation Assay

For pulldown and co-IP analysis of FER and LRR4, Agrobacteria containing *pFER::FER-GFP* (Escobar-Restrepo et al, 2007) and/or *p35S::LRR4-HA* (Ndinyanka et al., 2017) were infiltrated into *Nicotiana benthamiana* leaves. After 48 hrs of infiltration, the tobacco leaves were excised and grinded in liquid nitrogen. The tissue powder was re-suspended in extraction buffer [50 mM HEPES-KOH (pH 7.6), 150 mM NaCl, 1 mM DTT, 1 mM PMSF, protease inhibitor and 1 % NP-40]. The suspension was incubated on ice for 30 minutes and then centrifuged at 13,000 rpm for 20 minutes at 4°C. The supernatant obtained was then incubated with GFP-trap agarose beads or anti-HA agarose beads for 3-4 hours at 4°C on a rotating shaker. After incubation, the beads were washed three times with the extraction buffer containing 0.1 % NP-40 and boiled in SDS-PAGE loading buffer for 15 minutes at 75°C. The immunoprecipitates were then run on a 10% SDS-PAGE and transferred to nitrocellulose membrane to perform Western blotting.

## Acknowledgements

We are grateful to Y. Belkhadir, A.T. Fuglsang, N. Geldner, B. Gray, P.B. Larsen, G. Monshausen, and C. Zipfel for sharing published material; Elke Barbez for critical reading of the manuscript; Jit Thacker for help with preparing the manuscript; and the BOKU-VIBT Imaging Centre for access and expertise. This work was supported by the Austrian Academy of Sciences (ÖAW) (DOC fellowship to K.D.), Vienna Research Group (VRG) program of the Vienna Science and Technology Fund (WWTF), the Austrian Science Fund (FWF) (Projects: P26568-B16 and P26591-B16), the European Research Council (ERC) (Starting Grant 639478-AuxinER) (to J.K-V.), and the Swiss National Fund (SNF) (to C.R.).

## References

Barbez, E., Dünser, K., Gaidora, A., Lendl, T., & Busch, W. (2017). Auxin steers root cell expansion via apoplastic pH regulation in Arabidopsis thaliana. Proceedings of the National Academy of Sciences, 114(24), E4884–E4893.

Baumberger, N., Steiner, M., Ryser, U., Keller, B., & Ringli, C. (2003). Synergistic interaction of the two paralogous Arabidopsis genes LRX1 and LRX2 in cell wall formation during root hair development. The Plant Journal, 35(1), 71–81.

Braidwood, L., Breuer, C., & Sugimoto, K. (2013). My body is a cage: mechanisms and modulation of plant cell growth. New Phytologist, 201(2), 388–402.

Deslauriers, S. D., & Larsen, P. B. (2010). FERONIA is a key modulator of brassinosteroid and ethylene responsiveness in Arabidopsis hypocotyls. Molecular Plant, 3(3), 626–640.

Draeger, C., Fabrice, T. N., Gineau, E., Mouille, G., Kuhn, B. M., Moller, I.,… & Ringli, C. (2015). Arabidopsis leucine-rich repeat extensin (LRX) proteins modify cell wall composition and influence plant growth. BMC plant biology, 15(1), 155.

Duan, Q., Kita, D., Li, C., Cheung, A. Y., & Wu, H. M. (2010). FERONIA receptor-like kinase regulates RHO GTPase signaling of root hair development. Proceedings of the National Academy of Sciences, 107(41), 17821–17826.

Escobar-Restrepo, J. M., Huck, N., Kessler, S., Gagliardini, V., Gheyselinck, J., Yang, W. C., & Grossniklaus, U. (2007). The FERONIA receptor-like kinase mediates male-female interactions during pollen tube reception. Science, 317(5838), 656–660.

Fendrych, M., Leung, J., & Friml, J. (2016). TIR1/AFB-Aux/IAA auxin perception mediates rapid cell wall acidification and growth of Arabidopsis hypocotyls. Elife, 5, e19048.

Geldner, N., Dénervaud-Tendon, V., Hyman, D. L., Mayer, U., Stierhof, Y. D., & Chory, J. (2009). Rapid, combinatorial analysis of membrane compartments in intact plants with a multicolor marker set. The Plant Journal, 59(1), 169–178.

Gjetting, S. K., Ytting, C. K., Schulz, A., & Fuglsang, A. T. (2012). Live imaging of intra- and extracellular pH in plants using pHusion, a novel genetically encoded biosensor. Journal of experimental botany, 63(8), 3207–3218.

Humphrey, T. V., Bonetta, D. T., & Goring, D. R. (2007). Sentinels at the wall: cell wall receptors and sensors. New Phytologist, 176(1), 7–21.

Lewis, K. C., Selzer, T., Shahar, C., Udi, Y., Tworowski, D., & Sagi, I. (2008). Inhibition of pectin methyl esterase activity by green tea catechins. Phytochemistry, 69(14), 2586–2592.

Li, C., Wu, H. M., & Cheung, A. Y. (2016). FERONIA and her pals: functions and mechanisms. Plant physiology, 171(4), 2379–2392.

Löfke, C., Dünser, K., Scheuring, D., & Kleine-Vehn, J. (2015). Auxin regulates SNARE-dependent vacuolar morphology restricting cell size. Elife, 4, e05868.

Marre, E. (1979). Fusicoccin: a tool in plant physiology. Annual Review of Plant Physiology, 30(1), 273–288.

Ndinyanka Fabrice, T., Vogler, H., Draeger, C., Munglani, G., Gupta, S., Galatea Herger,… & Ringli, C. (2017) LRX Proteins play a crucial role in pollen grain and pollen tube cell wall development bioRxiv 223008; doi: https://doi.org/10.1101/223008

Ringli, C. (2010). The hydroxyproline - rich glycoprotein domain of the Arabidopsis LRX1 requires Tyr for function but not for insolubilization in the cell wall. The Plant Journal, 63(4), 662–669.

Scheuring, D., Löfke, C., Krüger, F., Kittelmann, M., Eisa, A., Hughes, L.,… & Kleine-Vehn, J. (2016). Actin-dependent vacuolar occupancy of the cell determines auxin-induced growth repression. Proceedings of the National Academy of Sciences, 113(2), 452–457.

Shih, H. W., Miller, N. D., Dai, C., Spalding, E. P., & Monshausen, G. B. (2014). The receptor-like kinase FERONIA is required for mechanical signal transduction in Arabidopsis seedlings. Current Biology, 24(16), 1887–1892.

Spartz, A. K., Ren, H., Park, M. Y., Grandt, K. N., Lee, S. H., Murphy, A. S.,… & Gray, W. M. (2014). SAUR inhibition of PP2C-D phosphatases activates plasma membrane H+- ATPases to promote cell expansion in Arabidopsis. The Plant Cell, 26(5), 2129–2142.

Spartz, A. K., Lor, V. S., Ren, H., Olszewski, N. E., Miller, N. D., Wu, G.,… & Gray, W. M. (2016). Constitutive expression of the auxin-related AtSAUR19 protein confers auxin-independent hypocotyl elongation. Plant Physiology, pp-01514.

Stegmann, M., Monaghan, J., Smakowska-Luzan, E., Rovenich, H., Lehner, A., Holton, N.,… & Zipfel, C. (2017). The receptor kinase FER is a RALF-regulated scaffold controlling plant immune signaling. Science, 355(6322), 287–289.

Wolf, S., Mravec, J., Greiner, S., Mouille, G., & Höfte, H. (2012). Plant cell wall homeostasis is mediated by brassinosteroid feedback signaling. Current Biology, 22(18), 1732–1737.

